# Microbial and Viral Genome and Proteome Nitrogen Demand Varies Across Multiple Spatial Scales Within a Marine Oxygen Minimum Zone

**DOI:** 10.1101/2022.11.11.516076

**Authors:** Daniel Muratore, Anthony D. Bertagnolli, Laura A. Bristow, Bo Thamdrup, Joshua S. Weitz, Frank J. Stewart

## Abstract

Nutrient availability can significantly influence microbial genomic and proteomic streamlining, for example by selecting for lower nitrogen to carbon ratios. Oligotrophic open ocean microbes have streamlined genomic nitrogen requirements relative to their counterparts in nutrient-rich coastal waters. However, steep gradients in nutrient availability occur at meter- and even micron-level spatial scales. It is unclear if such gradients also structure genomic and proteomic stoichiometry. Focusing on the eastern tropical North Pacific oxygen minimum zone (OMZ), we use comparative metagenomics to examine how nitrogen availability shapes microbial and viral genome properties along the vertical gradient across the OMZ and between two size fractions distinguishing free-living versus particle-associated microbes. We find a substantial increase in nitrogen content of encoded proteins in particle-associated over free-living bacteria and archaea across nitrogen availability regimes over depth. Within each size-fraction, we find that bacterial and viral genomic nitrogen tends to increase with increasing nitrate concentrations with depth. In contrast to cellular genes, the nitrogen content of virus proteins does not differ between size fractions. We identified arginine as a key amino acid in modulating the C:N ratio of core genes for bacteria, archaea, and viruses. Functional analysis reveals that particle-associated bacterial metagenomes are enriched for genes involved in arginine metabolism and organic nitrogen compound catabolism. Our results are consistent with nitrogen streamlining in both cellular and viral genomes on spatial scales of meters to microns. These effects are similar in magnitude to those previously reported across scales of thousands of kilometers.

**IMPORTANCE:** The genomes of marine microbes can be shaped by nutrient cycles, with ocean-scale gradients in nitrogen availability known to influence microbial amino acid usage. It is unclear, however, how genomic properties are shaped by nutrient changes over much smaller spatial scales, for example along the vertical transition into oxygen minimum zones (OMZs) or from the exterior to interior of detrital particles. Here, we measure protein nitrogen usage by marine bacteria, archaea, and viruses using metagenomes from the nitracline of the eastern tropical North Pacific OMZ including both particle-associated and non-associated biomass. Our results show higher genomic and proteomic nitrogen content in particle-associated microbes and at depths with higher nitrogen availability for cellular and viral genomes. This discovery suggests that stoichiometry influences microbial and viral evolution across multiple scales, including the micro- to millimeter scale associated with particle-associated versus free-living lifestyles.

## INTRODUCTION

A diverse consortium of marine microorganisms drive global biogeochemistry (1) and form the foundation of the marine food web. Major efforts have been undertaken to characterize the diversity and biogeographical patterns of marine microorganisms (2, 3) as well as their potential and realized metabolic output (4, 5). Emergent differences in the bioavailability of nutrients shaped by microbial activity, in turn, shape the evolution of microbial genomes and metabolism. Nutrient availability is an important mechanism in the evolution of gene content (6, 7), and has been implicated as the driver of observed patterns of genomic streamlining for bacterioplankton living in oligotrophic marine waters (8). Reduced nitrogen availability is thought to select for genomes with lower GC content and proteins with lower nitrogen to carbon ratios (9). More generally, the study of ‘stochio-genomics’ addresses how resource conditions can influence microbial genomic and proteomic elemental composition (7). For example, a survey of the North Pacific Subtropical Gyre at Station ALOHA found increasing genomic length, GC content, and nitrogen utilization in amino acid side chains from the mixed layer to the mesopelagic, corresponding to increasing nitrogen availability (10). Related work at Station ALOHA found metagenome-assembled genomes (MAGs) occurring on sinking particles collected at 150 meters were longer, had higher GC content, and predicted proteins had higher N usage than MAGs assembled from planktonic microbes at the same depth (11). Global marine metagenomes exhibit increased nitrogen content in arginine synthesis genes along the coasts as opposed to the open ocean, where nitrogen is presumed to be more limiting (6). On a global scale, marine metagenomic nitrogen content appears to correlate to nitrate concentrations (12). Studies linking nutrient availability to genomic streamlining have tended to focus on macro-scale patterns (e.g., spanning kilometers of depth changes or tens to thousands of kilometers between coast and open water) (7).

Steep chemical gradients can occur along much smaller spatial scales. Micronscale chemical heterogeneity imposes stark ecological transitions from the frame of reference of microorganisms (13). In marine environments, small particles such as sinking dust and phytoplankton cells constitute pockets of high availability and diversity of substrates for microbial metabolism (14). Taxonomic surveys of estuarine (15), coastal (16), and offshore (17) environments show that free-living microbial communities are taxonomically distinct from those on particles. Across these studies, particle-associated fractions are generally enriched in Gammaproteobacteria such as *Vibrio* and *Pseudoalteromonas*, while free-living communities contain more Alphaproteobacteria, such as the ubiquitous SAR11 group (15, 17). Free-living and particle-associated communities also differ functionally.

OMZs exhibit large differences in nutrient availability on small spatial scales - suggesting they are model ecosystems for exploring genomic and proteomic streamlin-ing. OMZs form in areas of enhanced nutrient loading, typically from upwelling (18). In these regions, nutrients fuel high surface primary production (19). Respiration of sinking organic matter by heterotrophs coupled with limited vertical mixing drives oxygen depletion at midwater depths, often to levels below detection (a few nM dissolved O2 (20, 21)). These dynamics result in stark vertical transitions between an oxic photic layer and a functionally anoxic, subphotic OMZ layer where nitrite accumulates as nitrate becomes the dominant oxidant for microbial respiration (22). The OMZ environment is ideal for testing stochio-genomic questions given that vertical gradients in bioavailable nitrogen and other key water chemistry parameters (e.g., total dissolved organic matter) are steeper in OMZ regions than in almost any other ocean environment (23, 24).

Despite evidence showing taxonomic and functional differences in particle-associated versus free-living marine microorganisms, such as the enrichment of copiotrophic gammaproteobacteria and increased prevalence of carbohydrate-degrading enzymes (2, 25, 11) there are limited data on how such differences affect stoichiogenomic properties (11). Based on prior research focused on macroscale spatial gradients, we hypothesize that stochiogenomic properties are also shaped by microscale resource variation between particle-associated versus non-associated niches. Specifically, we hypothesize that, compared to particle-associated microbes, free-living microbes exhibit genome-wide reductions in 1) GC content and 2) nitrogen content of encoded amino acids. We also hypothesize that these stoichiogenomic differences occur regardless of depth in the water column, i.e., that the genomic effects of resource heterogeneity (in this case, nitrogen) between particle and non-particle niches persist alongside changes in dissolved nitrate + nitrite concentrations with depth.

In this study, we analyzed 58 metagenomes sampled in 2013 and 2014 from the world’s largest OMZ, in the eastern tropical North Pacific (ETNP) (26). These metagenomes span the oxic surface, anoxic OMZ, and suboxic upper mesopelagic below the OMZ, and both particulate (> 1.6 micron) and free-living (0.2-1.6 micron) size fractions. We investigated drivers of metagenomic GC content as well as stoichiometric properties of bacterial, archaeal, and viral genes across depths and between size classes. We find trends in metagenome-level and gene-level GC content and amino acid nitrogen content that correspond to nitrate + nitrite concentrations similar to those seen in non-OMZ environments (10). We also find higher GC content and amino acid N content in particle-associated metagenomes for bacterial and archaeal genes. Viral genes do not exhibit stoichiogenomic differences between size fractions, although viral genes increase in nitrogen content with depth, similarly to bacteria. We selected an example core gene for bacteria, archaea, and bacteriophages and found patterns in amino acid composition that drive differences in bulk genetic and proteomic C:N ratios. For bacteria, functional analysis identified enrichment of genes involved in the synthesis of arginine, one of the amino acid drivers of bacterial core gene nitrogen content, on particle-associated samples where nitrogen remineralization and availability may be high. These results suggest heterogeneity in nitrogen availability on the microscopic scale in the vicinity of particles and on the meters scale of the oxycline in an OMZ can have as substantial an impact on microbial and viral genome evolution as does large-scale variability across ocean environments (e.g., pelagic vs. coastal) or ocean basins.

## RESULTS

### Biogeochemical Gradients in the ETNP OMZ

Metagenomic samples were taken from five stations in the region of the ETNP OMZ on two cruises in June 2013 and May 2014 (Figure 1a). Samples were serially filtered onto particle-associated (> 1.6 micron) and free-living (> 0.2 micron) size fractions across depths ranging from the surface to 2600m (see Methods). Oxygen saturation levels (see Methods) were between 95-101% for all stations in the surface mixed layer. The oxic-anoxic interface increases with depth offshore, but occurs in the top 100m of the water column with an average depth of approximately 80m (Figure 1 for representative profile). Dissolved nutrient profiles had canonical OMZ characteristics (22) (Figure 1). Using data from station 6 as an example (Figure 1), in the surface mixed layer nitrite concentrations were undetected and nitrate concentrations were below 1 uM. Between the base of the mixed layer (20m) and the oxic-anoxic interface (80m), nitrate concentrations increase to 20 *µ*M. Nitrite concentrations decline after peaking at 6.06 *µ*M at 125m, while nitrate concentrations continue to increase to above 40 *µ*M at 1000m.

**FIG 1.**
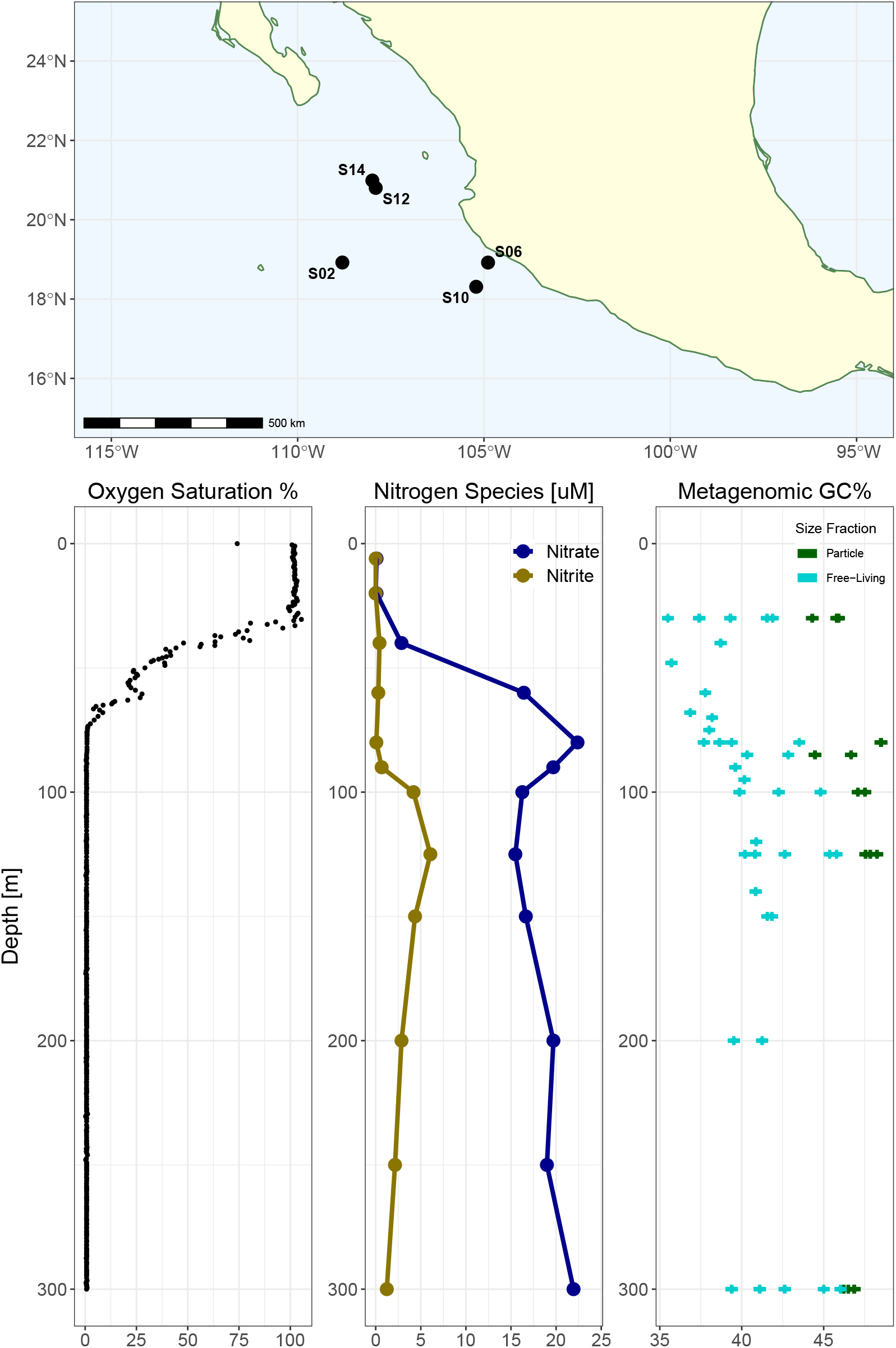
Environmental contextual data and sampling locations for this study. (a) Map showing sampling station locations for the 2013 and 2014 field collection efforts. (b) Representative oxygen saturation profile from Station 6 in the upper 300m. (c) Dissolved nitrate and nitrite profile for Station 6 measured in 2013. (d) Average GC content of metagenomic reads from all metagenomic samples taken from the upper 300m, representing a >1.6 micron particle-associated size fraction and a planktonic size fraction >0.2 microns.

### Genomic Streamlining Shaped by Nitrogen Availability at Small Spatial Scales

We analyzed the GC content of metagenomic reads to evaluate the potential for genomic streamlining given dissolved nitrate+nitrite differences across the water column and putative differences in nitrogen availability between size fractions. For both size fractions and across all samples, the average GC content of metagenomic reads (see Methods) is lowest at oxic surface depths, increases in alignment with nitrate concentrations in the anoxic OMZ core until approximately 150m, then remains roughly constant at lower depths (Figure 1d). While this trend was consistent between size fractions, particle-associated metagenomes had consistently higher GC content at all depths compared to free-living fraction metagenomes. Using a mixed effect model controlling for sequencing depth and spatial correlation in depth, we identified a significant effect of a 5.32% (± 0.587%) increase in metagenomic GC content between size fractions (see Methods, model parameters described in Supplemental Table 1).

We investigated 16S rRNA genes extracted from the metagenomes to explore the relationship between changing GC content and taxonomic composition (see Methods). Taxonomic patterns are summarized in Supplemental Figure 1 and match those reported in previous 16S-based surveys of the ETNP OMZ (27, 2). Key trends include a predominance of SAR11 Alphaproteobacteria at all depths throughout the water column, an abundance of nitrifying Thaumarchaeota along the oxycline, and an enrichment of Delta- and Gammaproteobacteria in particle-associated communities (Supplemental Figure 1). Notably, samples from the particle-associated fraction contained reads matching the mitochondrion of picoeukaryotes, primarily obligate endosymbiotic dinoflagellates from the Syndiniales group. The eukaryotic taxa we observed are consistent with 18S rRNA gene-based surveys in the ETNP OMZ (28).

### Spatial Structure in Stoichiogenomic Properties Vary Between Bacteria, Archaea, and Viruses

Next, we investigated the stoichiometric properties of nucleotide and amino acid sequences of microbial and viral genes throughout the water column and between size fractions. Metagenomes were assembled, and we generated a gene catalogue of 718,947 gene clusters from the assembled contigs (see Methods). We annotated 713,019/718,947 (99.1%) of the gene clusters using the KEGG orthology database (29). Annotated genes were divided between bacterial, archaeal, and viral genes for further analysis based on KEGG orthology search taxonomic assignments. We hypothesized that the nitrogen content of the amino acid sequences of predicted genes would follow trends similar to those of bulk metagenomic GC content. For each predicted gene, we calculated stoichiogenomic properties including GC content, the abundances of each amino acid in the encoded protein, the N:C ratio of amino acid side chains in the sequence, as well as codon usage bias (10). We generated ‘stoichiogenomic profiles’ for bacterial, archaeal, and viral gene datasets by a coverage-weighted average of these properties over all genes within a sample. For each dataset, we constructed an redundancy analysis (RDA) controlling for variable sequencing depth to identify ‘Domain’ (bacteria, archaea, or virus)-specific drivers of covariation in stoichiogenomic properties (see Figure 2). We modeled the relative explanatory effects of particle fraction and depth using a permutational analysis of variance (see Methods). The results of our ordination analysis are presented in Figure 2. The bacterial stoichiogenomic profiles (Figure 2a) split into two groups along the first RDA axis (64.46% of total variance). Positive values are associated with communities from the surface mixed layer (yellow points), and negative values are associated with OMZ and mesopelagic samples (blue points). The second axis (20.77%) separated free-living samples (triangles) with high values from particle-associated samples (circles) with lower values. The estimated *R*^2^ value for size fraction is 14.0% (*p* < 0.001) while the estimated *R*^2^ for sample depth is 22.1% (*p* < 0.001). For archaeal profiles (Figure 2b), the first axis (49.43% of variance) separated particle-associated samples at low values mostly from mesopelagic free-living samples at high values. The second axis (22.91%) places OMZ samples at higher values and surface mixed layer samples at lower values. The estimated *R*^2^ values for size-fraction and depth are 14.1% (*p* < 0.001) and 13.1% (*p* < 0.001), respectively. For viral profiles (Figure 2c), most of the variance is explained by the first axis (74.12%), which separates surface mixed layer communities from OMZ communities, with mesopelagic samples (dark blue) being in the middle. The *R*^2^ for size fraction for viruses is 1.2% (*p* = 0.363), while the *R*^2^ for depth is 29.5% (*p* < 0.001). While each domain has unique underlying variance structure in stoichiogenomic profiles, all three reveal patterns corresponding to water column depth, and in the case of Bacteria and Archaea, size fraction.

**FIG 2.**
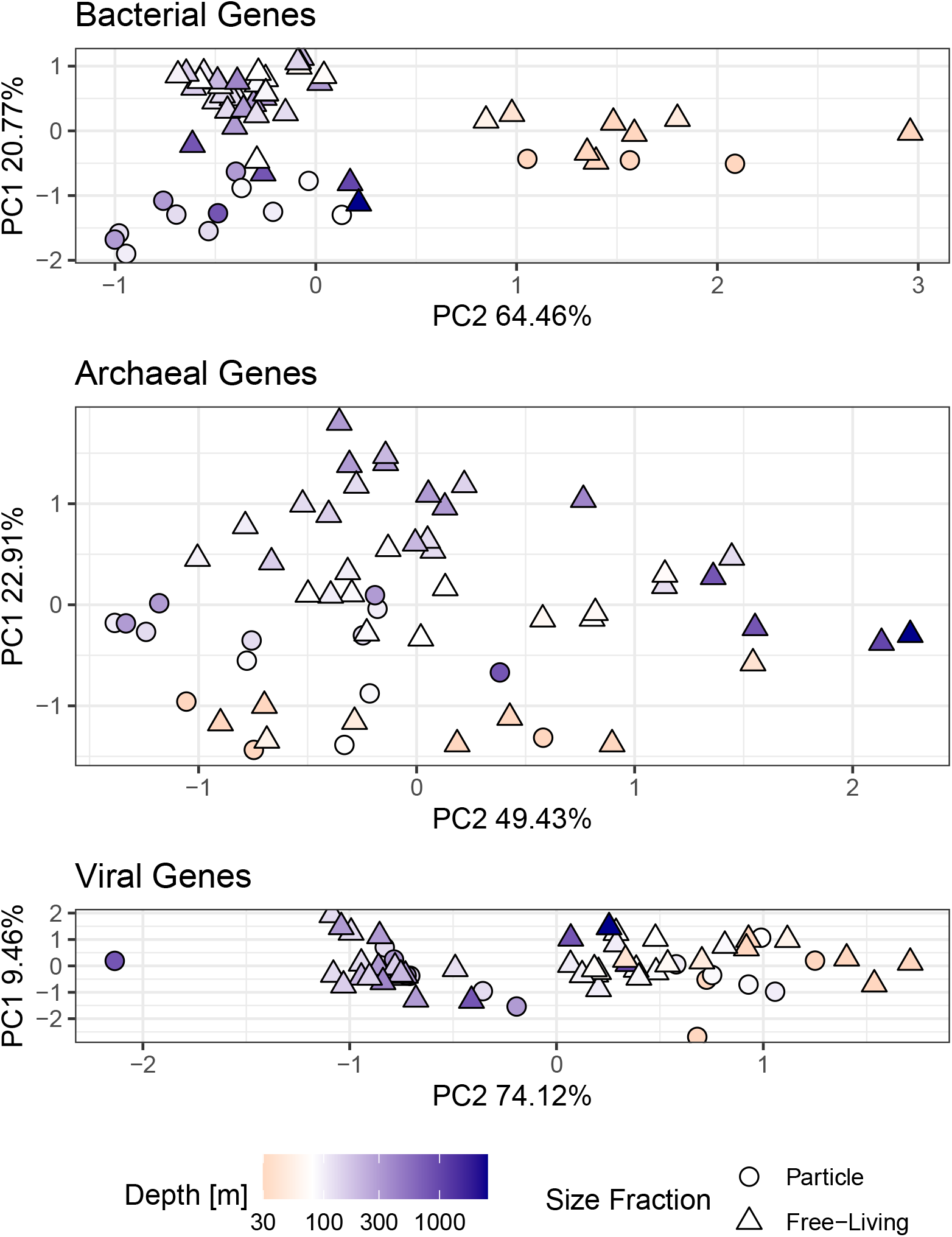
Stoichiogenomic profiles identify size-fraction and depth structure across Domains. Redundancy analysis (RDA) ordination was constrained by sequencing depth and the first two axes uncorrelated to sequencing depth are presented. Panels are separated by putative taxonomic assignment of the genes used to construct the ordinations. Point color indicates the sample depth, where yellow is above the average depth of the oxic-anoxic interface (80m) and blue is below the interface. Point shape indicates particle-associated (circle) versus free-living (triangle) samples.

### Particle Fraction Drives Stoichiogenomic Differences for Bacteria and Archaea

We constructed linear mixed-effect models of GC content, amino acid N:C ratio, and the number of nitrogen atoms in amino acid residue side chains (N-ARSC) to quantitatively test for differences between size fraction for each Domain (Methods). Our model structure included depth as a random effect to account for spatial autocorrelation and nonlinearity in the water column chemistry (Figure 1), so the depth effect cannot be expressed as a single parameter. Instead, we assessed the significance of depth by likelihood ratio tests against a null model without a depth-effect (Methods, estimated profiles in Supplemental Figure 2). For Bacteria, our models found significantly higher GC content (3.82% ±0.479%), NC ratios (0.00516±0.000406), and N-ARSC (0.0134±0.00102) in particle-associated communities (Supplemental Table 2, data plotted in Figure 3).We also found that depth had a significant effect for all three parameters, and the depth-effects increase GC and N-ARSC until reaching a maximum near 150m and remaining relatively constant with increasing depth, following the pattern of nitrate+nitrate (Supplemental Figure 2). For Bacteria, we also used the coverages of single copy core genes to estimate the average genome length for each sample (Methods) and found a significant increase in genome length (0.615 Mb ± 0.124 Mb, *p* < 1*e* − 5) in particle fraction samples (Figure 3).

**FIG 3.**
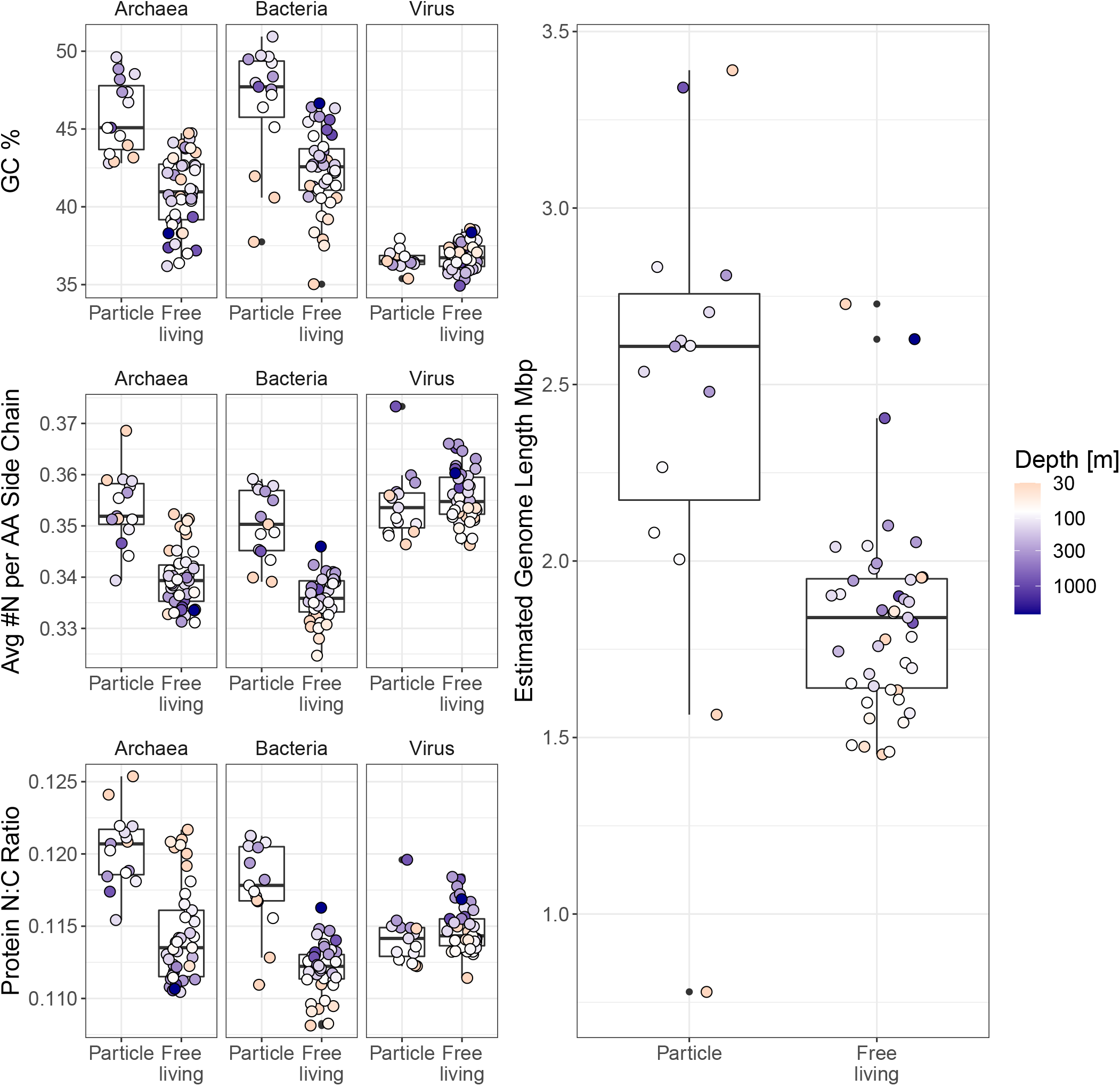
Microbial and viral genes exhibit stoichiogenomic structure across size fraction and depth. Each point represents a metagenome, where the value is the coverage-weighted average GC content (top row), nitrogen content of amino acid side chains (middle row), and total side chain N:C ratio (bottom row) of all archaeal, bacterial, or viral annotated genes in that metagenome. Box plots indicated the median, 25th and 75th percentiles of metagenomes within a size fraction. The right panel shows the distribution of estimated average bacterial genome length between size fractions as determined by the relative abundances of single copy core genes in each metagenome.

We found significant increases in archaeal GC content (4.58% ± 0.677%), NC ratios (0.00539 ± 0.000602), and N-ARSC (0.0128 ± 0.00152) in particle fraction samples (Supplemental Table 2). We also found a significant depth effect for N-ARSC and NC ratios for archaeal genes, but not for gene GC content. These effects are highest at the surface and decrease with depth, in contrast to the bacteria (Figure 3). For viruses, we did not find a significant increase in any stoichiometric parameters in the particle fraction. However, we did find a significant depth effect in NC ratios and N-ARSC for viral genes. Viral NC ratio and N-ARSC depth effects increased with depth similarly to bacteria (Figure 3, Supplemental Figure 2).

### Stoichiogenomic Differences are Characterized by Unique Amino Acid Frequencies

We chose one core functional gene for each Domain to explore whether metagenome-wide differences in stoichiogenomic properties were detectable on the single-gene level. Single gene-level analyses remove potential effects associated with differences in functional gene content between communities and focus on gene products that should have similar biochemical constraints to maintain the same function. We scanned the annotations of gene clusters for core functional genes that had many diverse representatives in the gene catalogue. There were 154 unique gene clusters annotated as bacterial *rpoZ*, the DNA-directed RNA polymerase omega subunit, 119 gene clusters annotated as archaeal *ftsZ*, the cell division protein, and 129 gene clusters annotated as phage structural protein Gp23. We used the coverage of the unique gene clusters in each sample to create a weighted average N:C ratio for *rpoZ, ftsZ*, and Gp23 for each sample. We then compared those weighted averages using the same mixed effect model framework to look for significant effects of particle size fraction on gene-specific N:C ratios. For bacterial *rpoZ*, we found an increase of 0.0304 (± 0.00891) in N:C ratio for particle-associated samples. For archaeal *ftsZ*, we found an increase of 0.00373 (± 0.00153). For viral Gp23, we found no significant difference in N:C ratio between size fractions. These single gene-level patterns match those of our model of genome-wide stoichiogenomic profiles (Supplemental Figure 3, model information presented in Supplemental Table 3).

We then investigated the amino acid composition of the gene clusters to understand the mechanisms governing their N:C ratios. We used the amino acid frequencies of each gene cluster to create a weighted average amino acid distribution for *rpoZ, ftsZ*, and Gp23 for each sample. We specified multiresponse linear models with a lasso penalty to learn the amino acids most predictive of size fraction, protein N:C ratio, and N-ARSC simultaneously (Methods). The parameters for these models are shown in Figure 4. Regularized regression allowed us to identify and rank the most important amino acids in predicting size fraction, protein N:C ratio, and N-ARSC for each Domain, excluding amino acids that do not contribute much information. The top driver of increased N:C ratios for all three Domains is arginine, the only amino acid with three additional N atoms on its side chain (Figure 4). High bacterial *rpoZ* N:C ratios are also driven by asparagine, lysine, and histidine. Archaeal *ftsZ* is more strongly associated with histidine than bacterial *rpoZ* and also is associated with increased tryptophan, unlike *rpoZ*. High viral Gp23 N:C ratios are also associated with increased histidine and asparagine, but are more strongly associated with glutamine than *ftsZ* or *rpoZ*. Interestingly, methionine is also associated with increased Gp23 N:C ratio, despite methionine having no N atoms on its side chain. This evidence shows that increased arginine is a common driver of stoichiogenomic variation across marker genes from all three Domains in our dataset, while there is variability in which other amino acids contribute to increased protein nitrogen content.

**FIG 4.**
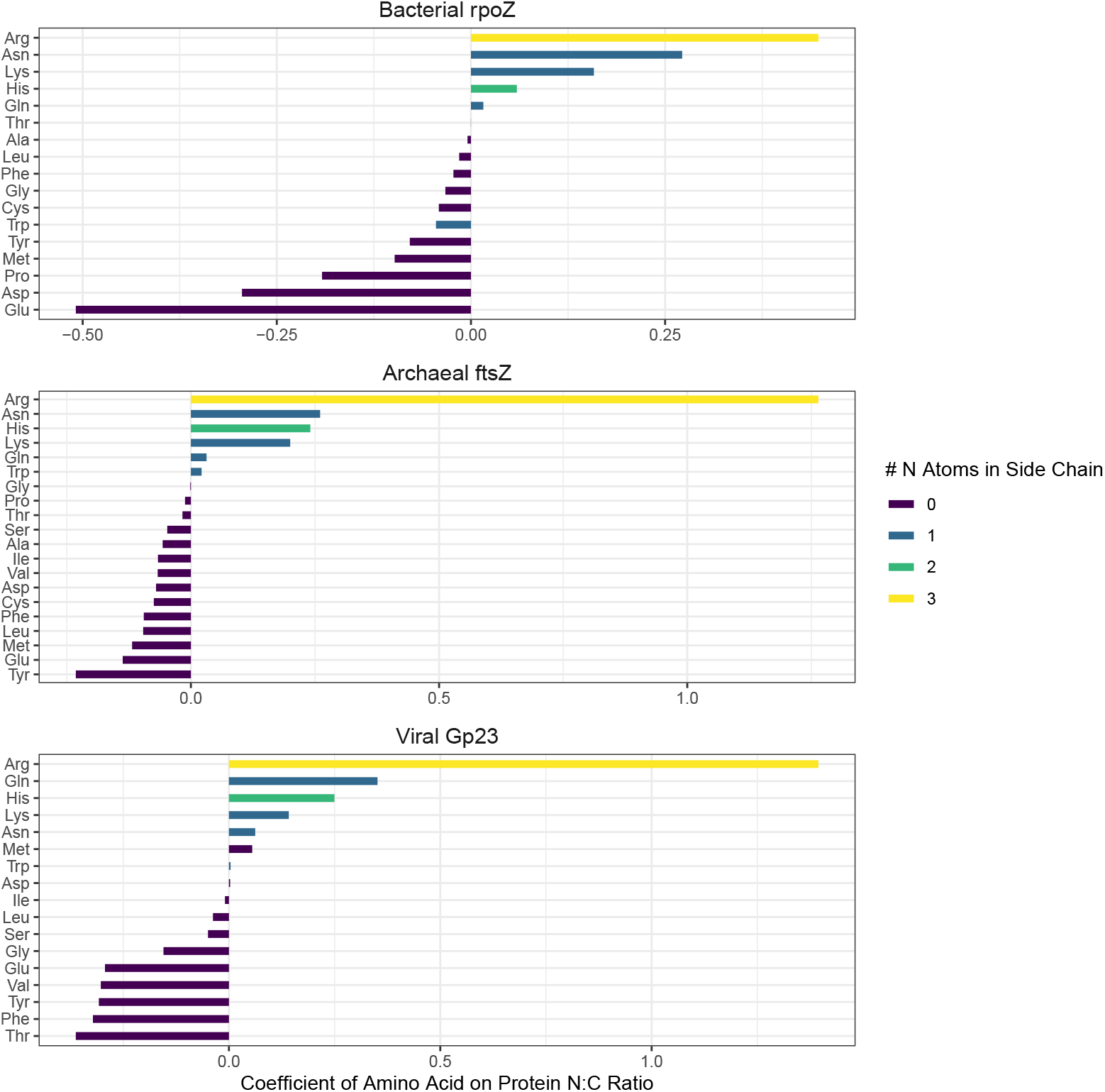
Domains have unique amino acid usage patterns to structure protein nitrogen content. Regularized amino acid regression parameters on average N:C ratios of widely conserved functional proteins across samples are presented. Positive values indicate increases with protein N:C ratio, and negative values indicate decreases. Amino acids are colored by the number of N atoms in their side chains. Note differences in scale between domains.

### Virus Structural Protein Amino Acid Frequencies Shift with Oxygen Concentration

Viral genes increased in GC, N-ARSC, and NC ratios with depth similarly to bacterial genes, although there was no significant differences for viral genes between size fractions for these parameters (Figure 3, Supplemental Table 2). While we did not find a significant difference between size fractions in the stoichiometry of a phage structural protein (Supplemental Table 4), we found that viral Gp23 and bacterial *rpoZ* N:C ratios both increase with increasing arginine, asparagine, histidine, lysine, and glutamine (Figure 4). We plotted depth profiles for those amino acids identified by our regression as contributing to Gp23 N:C ratios (Figure 5).

**FIG 5.**
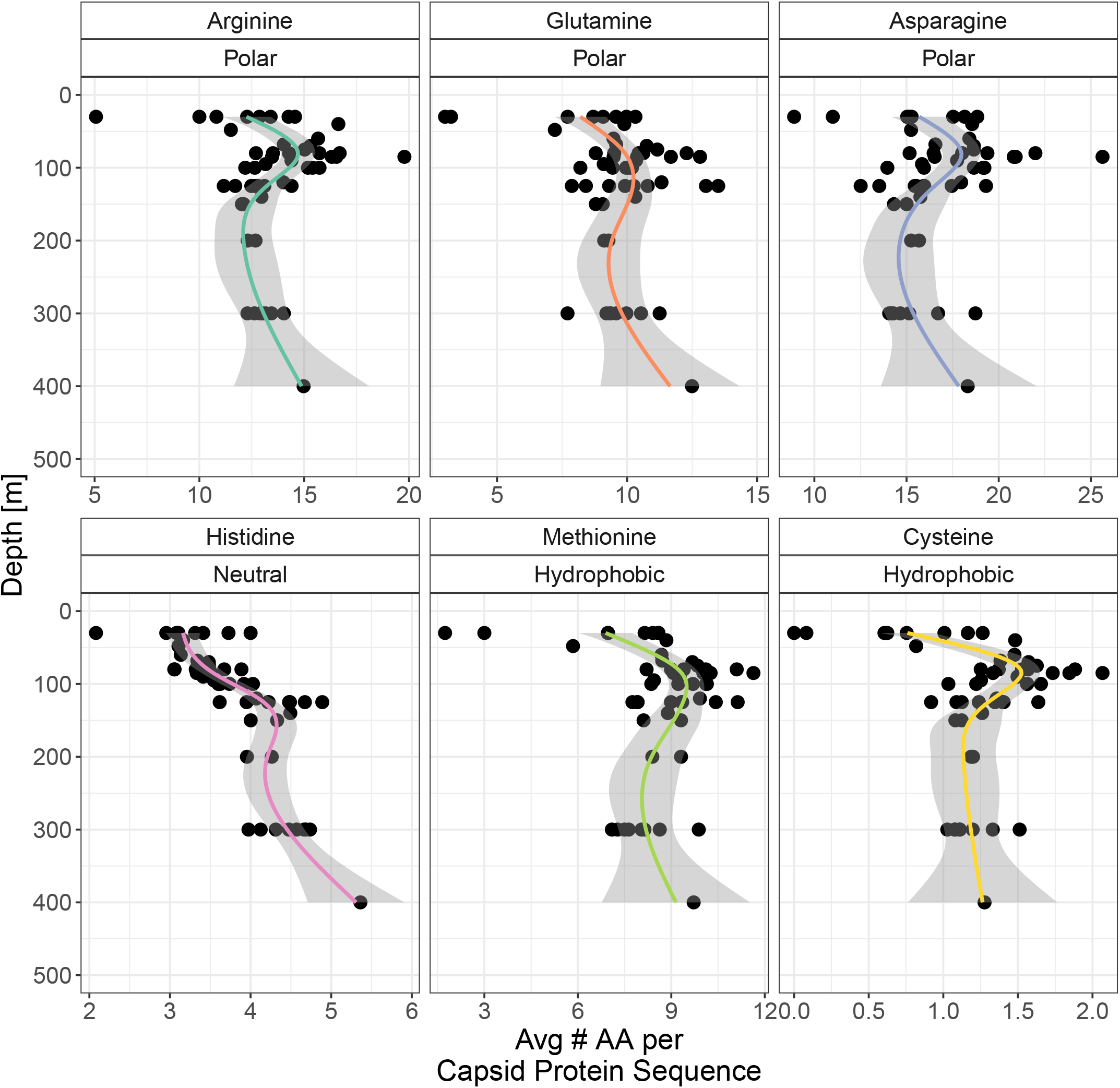
Viral Gp23 amino acid composition varies with depth. The x-axis represents the coverage-weighted average frequency of each amino acid in a Gp23 sequence for each sample. Amino acids are labeled as polar (hydrophilic)/neutral/hydrophobic. Smoothed lines to demonstrate overall trends are fitted via general additive model smoothing with shaded regions representing the 95% confidence intervals of the smoothing.

We plotted the profiles for the amino acids identified by the multiresponse regularized regression model (presented in Figure 4) to increase average Gp23 N:C ratios (Figure 5). Most of these amino acids have a peak in abundance slightly above 100m, which corresponds to the average oxic-anoxic interface depth across stations. Three of these amino acids had significant negative rank correlations with oxygen levels. Histidine (*ρ* = −0.487, *p* < 1*e* −5), an amino acid with two N atoms on its side chain, was negatively correlated with oxygen concentration, as was methionine (*ρ* = −0.311, *p* = 0.0334) (Figure 5). From our previous ordination of viral stoichiogenomic properties (Figure 2), we recalled that the primary axis clustered samples from depths around the oxic-anoxic interface at negative values, and samples from the surface mixed layer and mesopelagic samples at positive values. We then looked for other amino acids inversely correlated to oxygen concentration, indicating enrichment in high nitrogen anoxic water, and found cysteine (*ρ* = −0.228, *p* = 0.0334). Our amino acid-level analysis shows that the overall increase of viral N:C ratios with depth matches trends in bacteria and can be explained by a similar set of amino acids. Methionine and cysteine, the two hydrophobic and sulfur-containing amino acids, are specifically associated with the anoxic core of the OMZ and may contribute to the separation of that depth layer from mesopelagic waters in our earlier multivariate analysis of all viral genes.

### Particles are Enriched for Functional Genes Utilizing Organic Nitrogen Substrates

Our analysis of the stoichiometric properties of bacterial genomes indicates higher genomic and proteomic nitrogen content for particle-associated communities. This may be because some organic particles can act as either an organic nitrogen source directly, or a hotspot of nitrogen remineralization (30, 31). We compared the relative abundances of 412 bacterial KEGG orthologues between size fractions to identify which functional genes were enriched in particle fraction samples. We used a model based on the log-ratios of the relative abundances of each orthologue to a common ‘baseline’ orthologue to mitigate the negative constrained covariance and detection biases of compositional metagenomic data (32). Setting false discovery rate to 10%, we found 69 (16.8%) of KEGG orthologues enriched in particle-associated bacterial communities (Figure 6). Our analysis included 12 transporters, 5 of which take up branched chain amino acids or polyamines. The particle fraction was enriched in 4/12 transporters, including 3/5 transporters for organic nitrogen substrates. The particle fraction was also enriched in 9 genes related to amino acid degradation, including 4/13 of studied genes involved in the degradation of branched chain amino acids. The amino acid-level analysis identified arginine utilization as an important driver of protein-level N:C ratios for bacteria (Figure 4). The functional analysis also shows that 3/5 of studied genes in the arginine biosynthesis pathway are enriched in particle fraction bacteria (Figure 6). We also found enrichment on particle fractions for 3/4 of protein excretion genes and 3/4 of flagellum/pilus-related genes. For comparison, 8/30 of genes involved in carbon metabolism were enriched in the particle fraction. Our results suggest genes for uptake and degradation of organic nitrogen molecules are more abundant within particle-associated bacterial communities, consistent with the hypothesis that particles may offer usable organic nitrogen-containing substrates.

**FIG 6.**
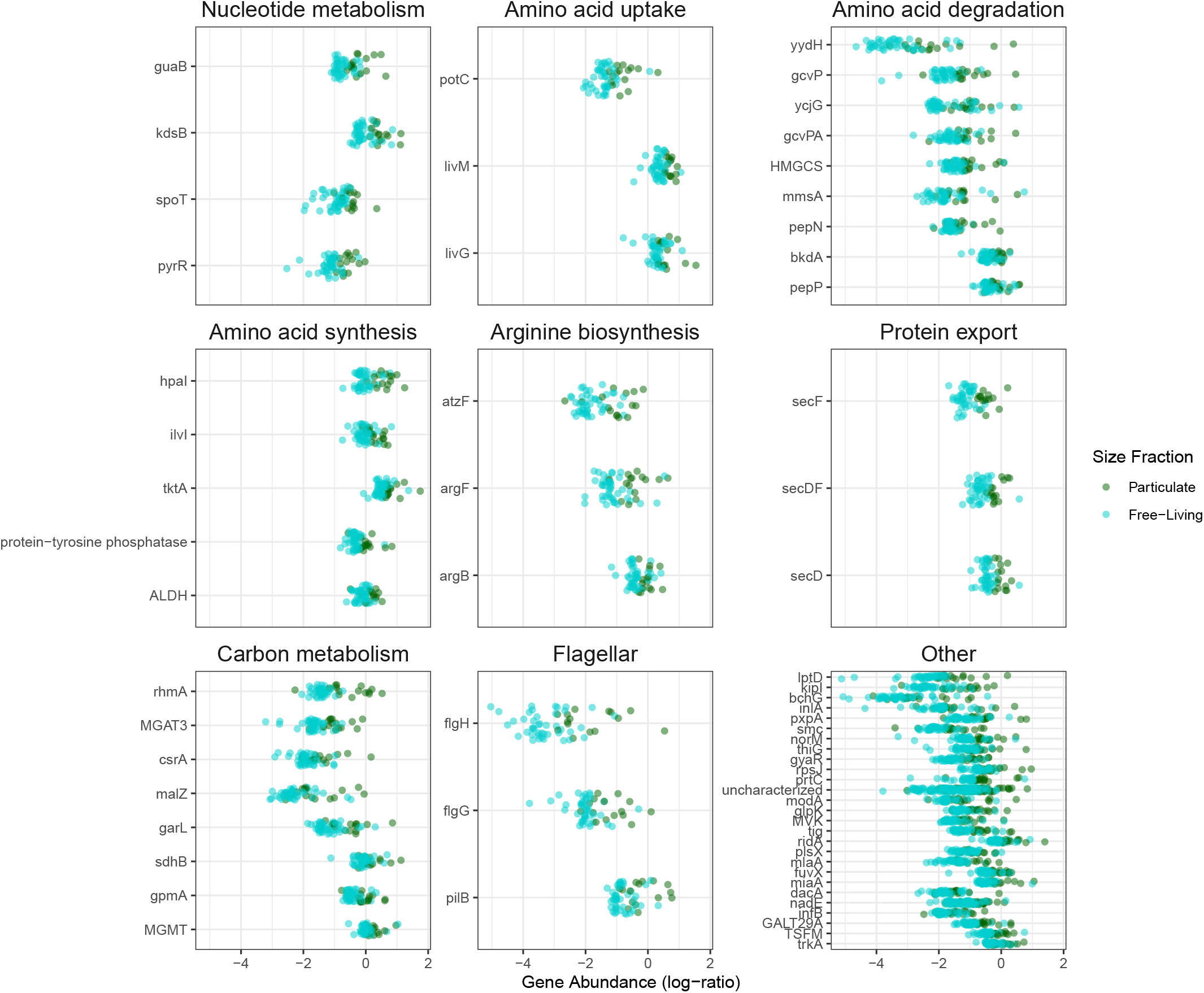
Bacterial functional gene content differences between size fractions. KEGG orthologues are grouped by metabolic pathways and identified by KEGG gene nomenclature (y axis). The log ratios of the relative abundance of each gene to *sdhC*, a core functional gene present across all samples, are plotted on the x-axis (see Methods). Point color indicates size fraction.

## DISCUSSION

Stoichiogenomic studies have identified genome streamlining-type evolutionary adaptations to nitrogen limitation across diverse marine microorganisms at spatial scales spanning kilometers of depth to thousands of kilometers across the ocean surface (7). Here, we examined the principles of stoichiogenomic streamlining at smaller spatial scales – across tens of meters surrounding the nitracline of the ETNP OMZ, and across microns separating planktonic and particle-associated microbes. The difference in nitrate+nitrite at the surface and at the oxic-anoxic interface (20 *µ*M over 60m) is equivalent to the difference in nitrate concentrations between the surface and 500m from a previous stoichiogenomic study in the North Pacific subtropical gyre (NPSG) (10), and mirrors average differences in nitrate+nitrite measurements between the surface and 350-400m in the Station ALOHA NPSG climatology [https://hahana.soest.hawaii.edu/hot/hot-dogs/]. 20 *µ*M is also similar to total nitrogen differences in surface concentrations between open ocean regions and high nitrate coastal upwelling regions (33). Our observed differences of nitrate+nitrite concentrations between the surface mixed layer and in the anoxic core is similar to the differences examined by previous stoichiogenomic studies at larger spatial scales. We found that predicted bacterial and viral protein nitrogen content correlated with the trend in nitrate+nitrite across depth. We also found that predicted bacterial and archaeal proteins are nitrogen enriched in particle-associated compared to planktonic metagenomes. For bacteria in particular, we could also link this gene-level variation to overall genome length as an additional parameter describing total genomic nitrogen quota. Furthermore, we found that the patterns across all genes hold for core marker genes, suggesting that stoichiogenomic differences between fractions and across depths are not driven exclusively by variation in community gene content.

Our analysis found that bacterial GC content and protein N-content were enriched on particles and increased with increasing depth and consequently increasing nitrate+nitrite. Similarly, bacterial genome lengths were longer in the particle fraction than in the planktonic fraction. Amino acid sequences for bacterial *rpoZ* in high nitrogen content samples had higher levels of arginine, asparagine, and lysine. Comparative functional analysis showed that arginine biosynthesis genes were more abundant in particle-associated communities, corresponding to the increased arginine frequency in *rpoZ* sequences. The enrichment in arginine biosynthesis genes in the particle fraction was accompanied by an enrichment in genes involved in the uptake and catabolism of other amino acids and organic nitrogen-containing compounds. Similarly, motility and secretory functions were enriched in the particle fraction. These functions have been previously correlated to particle-associated bacterial lifestyles (2).

Archaeal stoichiogenomic properties followed trends generally similar to those of bacteria, notably with nitrogen content enrichment in particle fraction samples. However, archaeal amino acid N content decreased with depth. Despite the trend having the opposite direction with depth, higher nitrogen archaeal sequences were associated with increased arginine, asparagine, and lysine, as in bacteria (Figure 4). Archaeal N:C ratios were also driven by histidine content (Figure 4). These results indicate a possibility that genomic streamlining may follow different evolutionary dynamics across different taxonomic groups, despite having a robust effect in overall genomic nitrogen content.

Beyond bacterial and archaeal populations, corresponding depth-related differences exist (although not differences between size fractions) in the nitrogen content of viral genes and protein sequences. Our analyses suggest stoichiogenomic adaptation to environmental nitrate+nitrite concentrations in viruses, adding to the body of stoichiogenomic literature thus far focusing on cellular life. Economizing viral elemental content to reflect the nitrogen limitation conditions of the host may be important for facilitating efficient infection (34, 35). Surveys of viral auxiliary metabolic genes in the eastern tropical South Pacific (ETSP) OMZ include viral genes for nitrogen cycling processes such as denitrification and assimilatory nitrate reduction (36), suggesting nitrogen metabolism influences viral adaptation in OMZ environments. We supplement this functional evidence by suggesting nitrate+nitrite concentrations may also correlate to the nitrogen quota of viral genomes. We also find stoichiogenomic parameters for viral genes in the anoxic core of the OMZ to be distinct from other depths via ordination analysis. This results contributes further evidence from an elemental composition and structural perspective to functional and taxonomic evidence showing unique viral communities in the eastern tropical South Pacific (ETSP) OMZ (36, 37, 38). Our evidence suggests that the uniqueness of OMZ viruses is not only present in taxonomy and functional gene content, but also in the genomic and amino acid chemical composition of viral particles.

Our amino acid-level analysis of the variability of a phage structural protein also identified elevated methionine and cysteine as a unique signature specific to the anoxic core. These amino acids are both hydrophobic and sulfur-containing. The increase of sulfur-containing amino acids in this layer may be related to complex sulfur cycling dynamics in OMZs (39). Hydrophobic residues, and specifically methionine, have been demonstrated to be important for phage capsid stability (40, 41). In the anoxic core of the ETNP OMZ, aerobic respiration rates are very low (19) and less efficient terminal oxygen acceptors are used (2). Furthermore, in the ETNP anoxic core, cell numbers decrease and the virus to microbe ratio increases (37). These factors suggest that encountering hosts with sufficient metabolic activity to support an infection may be more difficult at anoxic depths. Evidence exists for long-term (annual scale) persistence of viruses in the absence of hosts in the Red Sea (42), suggesting marine viruses may use persistence outside of hosts as an ecological strategy. The speculation that low metabolic rates may influence viral evolutionary strategies in the ETSP OMZ anoxic core is supported by genomic evidence that viruses have adaptations for slow and intermittent replication (37). Replacement of structural protein amino acid residues with stabilizing hydrophobic residues, such as the cysteine and methionine enrichments we find, may be a novel mechanism aiding in viral structural stability, allowing for longer residence times between host encounters to overcome low host abundance and metabolic rates.

Despite finding viral genomic streamlining with depth, we were unable to detect a significant difference in the stoichiogenomic properties of viral genes between particle-associated and planktonic metagenomes. This leads us to at least two hypotheses – (i) selective forces for streamlining differ between planktonic viral and particle-associated viral communities and are potentially weaker for viruses that infect hosts that exhibit planktonic and particle-associated lifestyles; (ii) viruses detected on particles do not necessarily infect particle-associated hosts and instead passively aggregate onto particles. To the latter hypothesis, recent molecular studies of particulate metagenomic communities have recovered cyanophage DNA in abyssal depths (43), suggesting that viral DNA can potentially be vertically transported on particles and persist where hosts may not be metabolically active. Our finding that the stoichiometric properties of planktonic and particle-associated viral genes are similar lend further evidence to the idea that viruses with planktonic-adapted hosts may passively aggregate to particles.

In summary, this study demonstrates that stoichiogenomic differences occur on spatial scales much smaller that spatial scales previously studied (6, 10). We show genomic differences in GC content of 3-5% between particle-associated and free-living metagenomes from the same depth, similar to previously reported differences observed over ranges of hundreds of meters from Mende et al. (10). The increase in N-ARSC observed in particle-associated versus free-living metagenomes is within the range (4-7% difference) between coastal and open ocean proteins (6). We also offer evidence of the extension genomic streamlining with nitrate+nitrite concentrations from the surface mixed layer to the OMZ anoxic core (30-100m) to viruses, despite not having their own independent metabolism or nutrient uptake. Better understanding the factors shaping the nutrient requirements of marine microbes (and their viruses) is vital to understanding the mechanisms shaping marine ecology and biogeochemistry. Macromolecules with compositions susceptible to stoichiometric forcing (e.g., proteins and genomes) can comprise a substantial proportion of cell contents for small cells (44). Therefore, evolutionary flexibility in these traits may have important emergent biogeochemical effects at larger scales. Viral infection can also shape the ecology and biogeochemistry of microbial communities (35). The elemental composition of viruses can generate lysate with distinct elemental ratios compared to the host (45). Understanding the influence of viral infection on the stoichiometry of marine detrital organic matter, and therefore carbon cycling, requires understanding the evolutionary relationships between host and viral elemental composition (46). Here we show evidence of viral genomic streamlining occurring in parallel to hosts along nitrate+nitrite concentration gradients with depth. Including stoichiogenomic effects, such as those described in this study, in future ecosystem models may offer an important route to our future understanding of the role of microbial adaptation and macromolecular contents in biogeochemical cycling.

## Author Warranty

If accepted for publication, the Work will be made freely available to the public on ASM’s *mSystems* website. ASM will grant the public the nonexclusive right to copy, distribute, adapt, and transmit the published Work for commercial or non-commercial use with proper attribution under the Creative Commons, Attribution license, Version 4.0 (CC-BY). For details, see https://creativecommons.org/licenses/by/4.0/ and https://creativecommons.org/licenses/by/4.0/legalcode,as well as https://journals.asm.org/author-warranty-and-provisional-license-publish.

## MATERIALS AND METHODS

### Metagenomic Data

All metagenomic data from the ETNP were sequenced on the Illumina platform and are associated with BioProject ID PRJNA632347.

### CTD and Nutrient Data

A Sea-Bird conductivity, temperature, and depth (CTD) sensors were deployed with Niskin rosettes during sample collection on a package including a fluorimeter, transmissometer, and Sea-Bird SBE43 oxygen sensor (47). Samples for nitrate and nitrite nutrient analyses were collected and 2013 samples were processed as described in Ganesh et al (2) and Glass et al. (47). Samples for nitrate and nitrite analysis for the 2014 cruise were processed as described in Ganesh et al. (48). For this study, down-casts of CTD profiles were analyzed using TEOS’s python Gibbs Seawater (gsw) toolbox v. 3.4.0 loaded into R via the reticulate library v. 3.38.0. Using latitude, longitude, pressure, temperature, and salinity readings, corrected height relative to sea level was calculated along with absolute salinity and conservative temperature. Absolute salinity and conservative temperature were used to derive density anomaly sigma0 with a reference pressure of 0 decibar. These properties were also used to calculate oxygen solubility, and oxygen saturation was derived from the oxygen solubility and oxygen readings from the CTD instrumentation package. Data were then smoothed by binned averaging to the nearest 1m of height.

### Sequence Data Quality Control, Taxonomic Assessment, and Assembly

Interleaved reads were deinterleaved using Nathan Haigh’s deinterleaving script from github https://gist.github.com/nathanhaigh/3521724. Reads for all metagenomes were then adapter trimmed and quality filtered using TrimGalore https://www.bioinformatics.babraham.ac.uk/projects/trim_galore/ with a Phred quality cutoff score of 25 and a minimum read length of 100bp. After trimming and filtering, reads were merged using FLASH (49) with expected read length adjusted to 200bp to minimize exclusion of mostly overlapping reads.

Before paired-end reads were merged, putative small subunit (SSU) ribosomal reads were identified using Metaxa2 (50). All identified 16S reads were then extracted from the forward and reverse reads for each metagenome and the sequences were quality assessed and dereplicated using dada2 as implemented in R (51). Sequence variants were then assigned taxonomy using the SILVA v1.3.2 database (52) from dada2’s assignTaxonomy function.

Parameters on the distributions of merged paired-end reads (average length, mean and variance of GC content, total merged reads) were calculated for each metagenome using Biopython utilities. We modeled the average GC content of metagenomic reads for each sample using a mixed-effects linear model with log10 total number of reads and size fraction as fixed effects and depth as a random effect using exponential correlation structure to account for spatial autocorrelation in the water column.

Metagenomes were assembled using megahit (53) with a starting kmer length of 27 and default parameters. Contig statistics were then calculated using custom python and R scripts.

### Gene Catalogue

Using BioPython (54) utilities, assemblies were filtered for all contigs of length > 1000. Then, genes were predicted for every metagenome using Prodigal v. 2.6.3 with standard parameters on the metagenome setting (55). Partial genes were removed.

After combining all predicted genes from all metagenomes, a nonredundant catalog of genes was creating using CD-HIT with amino acid sequences, clustering genes with greater than 95% identity and 90% alignment of the shorter sequence (56). To identify COGs represented in the clusters, FetchMG’s COG extraction script was used with standard parameters (57).

### Calculating Gene Characteristics

Using functionalities from scripts provided in Mende et al. (10) as well as additional BioPython utilities, gene length, gene GC content, molecular weight of summed amino acids coded by each gene, codon usage (as defined in Mende et al, a ranking scheme for determining the diversity of codons used for a particular amino acid in a given gene), codon bias, the number of nitrogen and carbon atoms in residue side chains for all coded amino acids (N/C-ARSC), NC-ratios of those side chains, as well as individual amino acid counts were calculated for each gene. Additionally, the bulk GC content of each metagenome was calculated for all merged reads using Biopython.

### Read Mapping for Abundance and Genome Size Estimation

The BWA-MEM algorithm was used to map individual metagenomes to the assembled gene catalog using standard parameters (58). After mapping, results were filtered for only alignments with 95% or higher identity (calculated as 1-(number of mismatches)/(alignment length)) and an alignment lengths greater than 60 bp, using a mix of samtools (59) and the pysam python package (https://www.osti.gov//servlets/purl/1559931). Resulting alignments were then converted into coverage for each gene, dividing the total number of base pairs mapped to each reference gene by the gene length. To estimate average genome copy number, these coverages were normalized to the average coverage of ten single-copy COGs (60), the same method used in (10). Average genes per genome were then calculated by summing the copy numbers for all genes for each metagenome using a custom script in R. Average weighted gene coverage was then calculated by dividing the coverages for each gene in a metagenome by the sum coverage of all genes in that metagenome, to mitigate the differences of sequencing depth between samples.

To translate gene properties to a metagenome-wide summary, the gene properties for each gene were then multiplied by their weighted coverage and summed for each metagenome. This created a ‘gene property profile’ for each metagenome which was then used in further statistical analyses.

### Statistical Analyses

Ordination Analysis Because our samples were sequenced across a wide range of depths, and Illumina sequencing has been shown to display molecular bias in sequencing (61, 62), we aimed to carefully undergo statistical analyses keeping this effect in mind. Therefore, for ordination analyses, we conducted RDA using sequencing depth as a covariate, then removing that axis from the ordination to ensure all patterns in underlying data structure were uncorrelated to sequencing depth. For all regression models, sequencing depth was included as an independent variable to reduce the potential impact of confounding on parameter estimation.

To first explore differences in covariation between metagenomic gene properties, an RDA was constructed using the vegan package v. 2.5-7 in R (63) based on all properties listed in Calculation of Gene Characteristics. Two axes were found to be associated with eigenvalues larger than the eigenvalue associated with sequencing depth, so they were selected for presentation. Post-hoc permutational analysis of variance tests were conducted to find the percent variance explained by size fraction and asinh-transformed depth for each group independently. We transformed the depth in order to account for nonlinearity in the water-column chemical structure, which changes more gradually with increasing depth below the anoxic core of the OMZ.

Modeling Stoichiogenomic Parameters We structured the comparison between size-fractions for stoichiogenomic parameters among bacteria, archaea, and viruses as separate linear mixed effect models. We specified depth as a random effect with exponential correlation structure to represent spatial autocorrelation in water column chemistry, and to capture nonlinear relationships between depth and stoichiogenomic parameters. Sequencing depth was also used as a covariate in this model to control for biases described above. Model parameters were learned using maximum likelihood methods via R’s nlme package v 3.1-155. For each model, another model was learned using only fixed effects and not including depth to assess the significance of including depth in the model. Model comparisons were conducted via likelihood ratio tests and comparison of Akaike Information Criterion (AIC) to control for the difference in model complexity. For models where there was a significant likelihood gain in including depth, the random effect estimate learned for each depth was correlated to depth using spearman’s *ρ*.

Amino Acid Analysis First, the coverages of all KEGG orthologues and the number of gene cluster representatives in the gene catalogue were assessed. For each domain, we selected the orthologue with the highest total coverage, coverage in all samples, the greatest number of gene cluster representatives in the gene catalogue, and a general function presumed to be universal to diverse taxa within each domain. These genes, *rpoZ* for bacteria, *ftsZ* for archaea, and Gp23 for viruses, were considered our ‘marker genes’ for amino acid frequency analysis. The amino acid counts for the representative sequence of each gene cluster for each marker gene were then used to construct a weighted average amino acid profile for each sample, using the coverages of each gene cluster in that sample as the weight.

The weighted amino acid averages for each Domain were centered and scaled to account for different dynamic ranges. We generated one model for each Domain. The models used the scaled amino acid ranges as predictors and the sample size fraction (binarized as particle fraction=1, planktonic=0), scaled weighted average N-ARSC for the Domain protein, and scaled weighted average N:C ratio of the Domain protein as multiresponses. For example, for bacteria, we used the average number of alanine, cysteine, etc., for each *rpoZ* in each sample, weighted by the abundances of each *rpoZ* sequence, to simultaneously predict the size fraction, average N-ARSC of *rpoZ* sequences, and average N:C ratio of *rpoZ* sequences for that sample. We learned the parameters of a linear model using a maximum likelihood framework with an *L*_1_ regularization to reduce redundancy among amino acids. 70% of metagenomes were used for model learning and cross validation, while 30% were used for testing. Models were learned using the R glmnet package 4.13 ‘multigaussian’ family of models. Cross validation using 10 folds was conducted to optimize the regularization parameter, and the model with the minimum average mean squared error across all 10 folds of the test data was selected. The model was fitted and then assessed against the remaining 30% of test data (See Supplemental Figure 4 for model evaluation).

Functional Analysis A log-ratio approach was used to compare the relative abundances of functional genes between samples. Log-ratios have been recommended specifically for use in metagenomic datasets to account for sequencing bias and to overcome negative constrained covariance structure surrounding compositional data (32).

In order to construct log-ratios, we selected *sdhC*, one of the components of succinate dehydrogenase from the TCA cycle, as a focal gene to be the denominator. This gene is part of a central metabolic pathway widely distributed across bacteria, and had among the highest coverage in all of our samples. This left 412 remaining KEGG orthologues that had at least some reads mapping in all samples. For each orthologue, we summed the coverages of all gene clusters annotated as that orthologue for each sample, divided it by the summed coverage of *sdhC* in that sample, and then took the log to reduce variance. Then, a linear model was constructed for each orthologue using the log-ratio as the response, and size fraction, depth, and a size-fraction/depth interaction term as predictors. The estimates for all three of these parameters were retained, and then an FDR threshold at 10% was applied to the p-values of all parameters across all models to account for multiple testing.

### Data Availability

All metagenomic data from the ETNP were sequenced on the Illumina platform and are associated with BioProject ID PRJNA632347. Intermediate and final data products and code are available in repository https://github.com/dmuratore/omz_stoichiometry with archival hosting at DOI: https://zenodo.org/badge/latestdoi/560085581.

## SUPPLEMENTAL MATERIAL

This manuscript references supplementary materials, as described below.

**FIG S1**. 16S marker gene community composition of OMZ metagenomes. Total numbers of 16S reads extracted from metagenomes (Methods) are coded by taxonomic affiliation. PF indicates particle fraction, SV indicates planktonic fraction. Stations are coded by S (station number), and the depth in meters is shown in the y axis.

**FIG S2**. Random effects of depth on stoichiogenomic parameters. Random effects are shown for parameters where including a random depth effect resulted in significant gain in model likelihood (likelihood ratio test, p<0.05). Each point represents the effect on the parameter associated with that sample depth. A general additive model smoothing is applied to show the relationship with depth, shaded area represents 95% confidence intervals on the smoothing.

**FIG S3**. N:C ratios of marker gene sequences. Points are colored by size fraction, and general additive model smoothing with 95% confidence intervals (shaded area) are overlaid.

**FIG S4**. Model evaluation for amino acid regression. Plots of predicted (x) and actual (y) values for test data are presented with the 1:1 line for each Domain. Model parameters reported are the slope and adjusted *R*^2^ for the linear regression of predicted to actual.

**TABLE S1**. Mixed Effect Model Statistics for Bulk Metagenomic GC Content. A mixed effects model was used to estimate the difference in total average GC content of metagenomic reads between size fractions. The correlation length scale of the exponentially distributed random effect due to sample depth is reported along with estimates of the fixed effects for size fraction and sequencing depth on GC content.

**TABLE S2**. Mixed Effect Model Statistics for Testing Significant Effect of Particle Fraction on Stoichiogenomic Parameters. Models were constructed for each stoichiometric parameter for each Domain separately. Estimates and standard errors for the difference in means between particle and free-living fractions are presented with p-value, as well as total model AIC and log-likelihood for the mixed effect model with a random depth effect. The likelihood ratio compares a model with the random depth effect to a model without a depth effect, and a p-value for the corresponding log-ratio test. For models with a significant depth effect (likelihood ratio test p<0.05), the Spear-man rank correlation of the random depth effect with depth is presented with a p-value. A positive value means in general the random effect increases with increasing depth, a negative value means the effect decreases with increasing depth.

**TABLE S3**. Model Parameters for Multiresponse LASSO Regression Models. Table contains information for the parameter value in estimating protein N:C ratio for bacterial *rpoZ*, archaeal *ftsZ*, and viral Gp23, as well as the N-ARSC value of the amino acid.

**TABLE S4**. Model Statistics for Core Gene Mixed Effect Models. Mixed effect model statistics for model setups matching those presented in supplemental table 2, but for bacterial *rpoZ*, archaeal *ftsZ*, and viral Gp23 individually are presented.

## ACKNOWLEDGMENTS

The authors thank the captain and crew of the R/V Oceanus for enabling sample collection. This work was supported by National Science Foundation grants 1151698, 1558916, 1564559, 2130185 to FJS, Danmarks Frie Forskningsford 9040-00327B to LAB, Simons Foundation grant 721231 to JSW. Funding agencies had no role in conducting the research. DM conceptualized, performed bioinformatic analysis, performed statistical analysis, and wrote the manuscript. ADB and FJS collected samples, conceptualized, generated sequence data, and advised in analysis, writing, and editing of the manuscript. LAB and BT carried out fieldwork, processed samples for nutrient analysis, and advised in writing and editing the manuscript. JSW contributed to development of analytic methods, analysis, and editing of the manuscript.

**Please read the Instructions to Authors or browse the FAQs for further details**.

