## Supplemental Materials for "Microbial and Viral Genome and Proteome Nitrogen Demand Varies Across Multiple Spatial Scales Within a Marine Oxygen Minimum Zone"

Daniel Muratore,[a,b,c] Anthony D. Bertagnolli,[d] Laura A. Bristow, [e] Bo Thamdrup,[e] Joshua S. Weitz,[b,f,g] Frank J. Stewart [d,h,*]
Georgia Institute of Technology, Interdisciplinary Program in Quantitative Biosciences, School of Biological Sciences, Atlanta, United States [a]
Georgia Institute of Technology, School of Biological Sciences, Atlanta, United States [b]
Santa Fe Institute, Santa Fe, United States [c]
Montana State University, Department of Microbiology and Cell Biology, Bozeman, United States [d]
University of Southern Denmark, Department of Biology, Odense, Denmark [e]
Georgia Institute of Technology, School of Physics, Atlanta, United States [f]
École Normale Supérieure, Institut de Biologie, Paris, France [g]
Georgia Institute of Technology, Center for Microbial Dynamics and Infection, Atlanta, United States [h]

Please send correspondence to:
Daniel Muratore,

This file includes:
Supplemental Figures S1-S4
Supplemental Tables S1-S4

**Figure S1.** 16S marker gene community composition of OMZ metagenomes. Total numbers of 16S reads extracted from metagenomes (Methods) are coded by taxonomic affiliation. PF indicates particle fraction, SV indicates planktonic fraction. Stations are coded by S (station number), and the depth in meters is shown in the y axis.

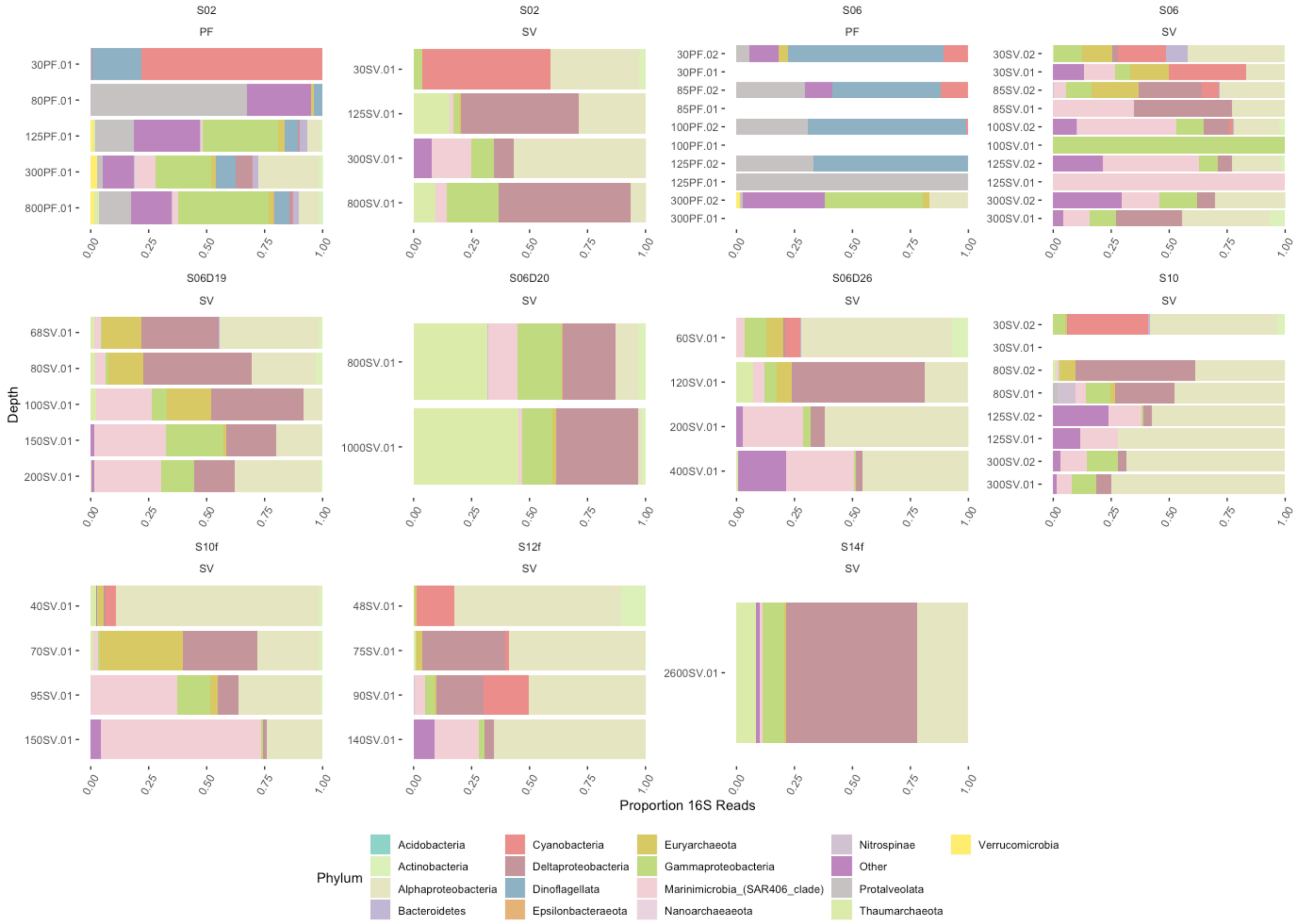

**Figure S2.** Random effects of depth on stoichiogenomic parameters. Random effects are shown for parameters where including a random depth effect resulted in significant gain in model likelihood (likelihood ratio test, $p<0.05$). Each point represents the effect on the parameter associated with that sample depth. A general additive model smoothing is applied to show the relationship with depth, shaded area represents 95% confidence intervals on the smoothing.

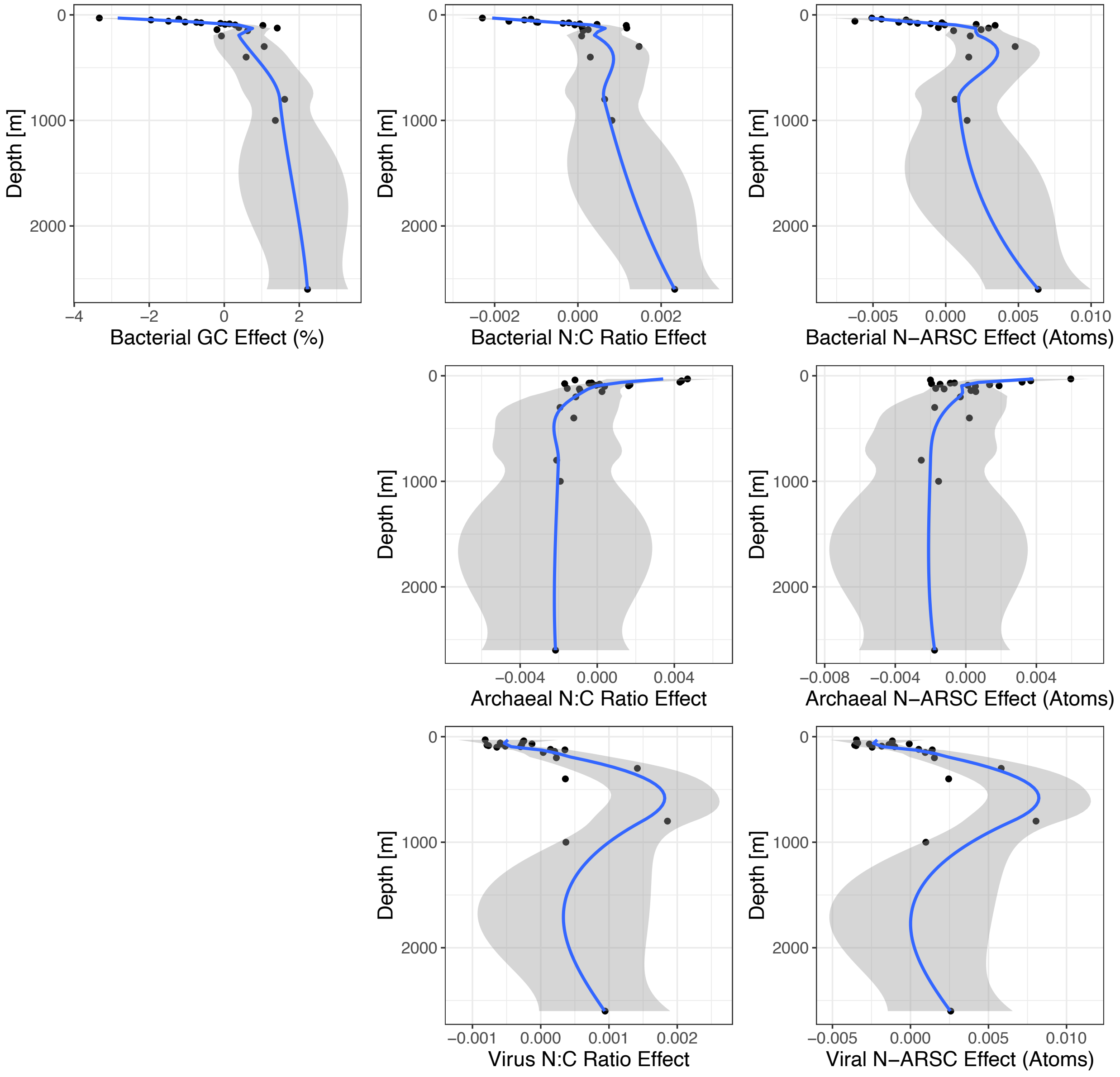

**Figure S3.** N:C ratios of marker gene sequences. Points are colored by size fraction, and general additive model smoothing with 95% confidence intervals (shaded area) are overlaid.

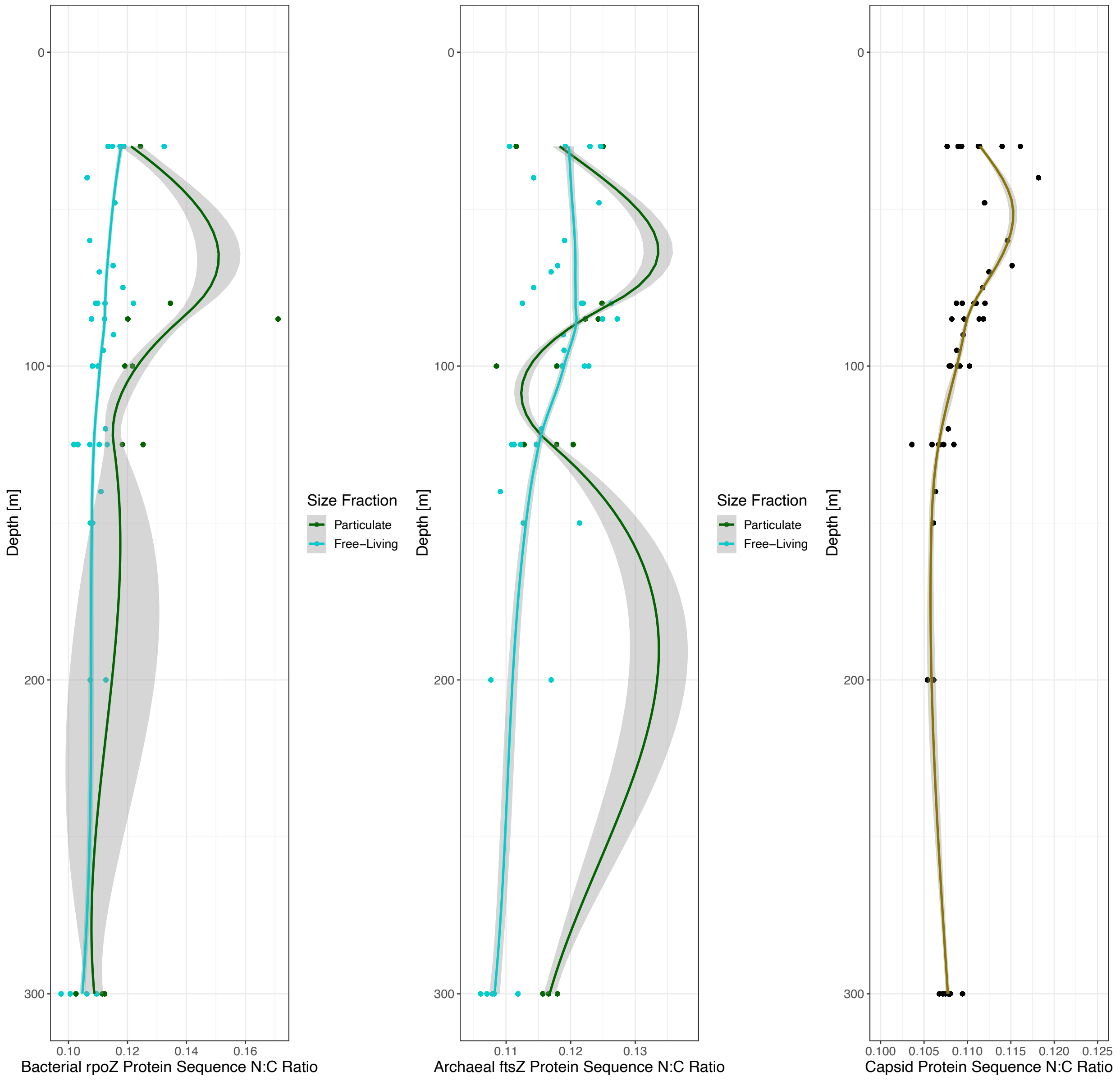

**Figure S4.** Model evaluation for amino acid regression. Plots of predicted (x) and actual (y) values for test data are presented with the 1:1 line for each Domain. Model parameters reported are the slope and adjusted $R^2$ for the linear regression of predicted to actual.

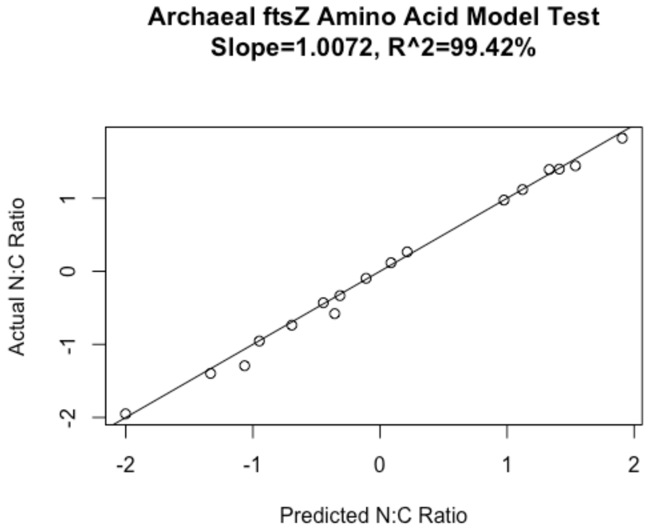


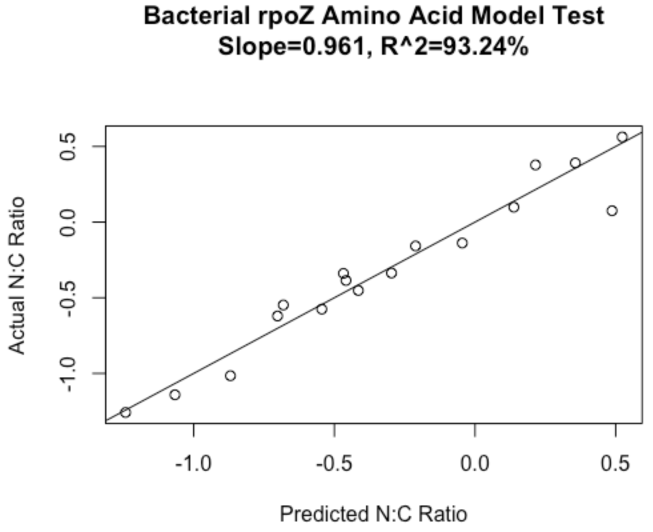

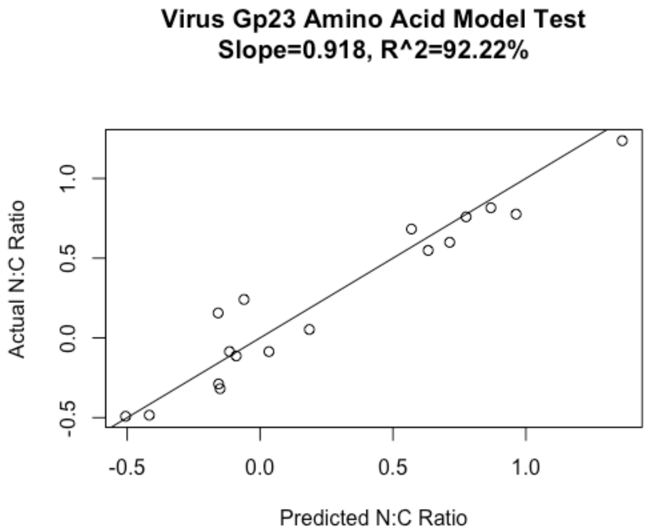
Virus Gp23 Amino Acid Model Test
Slope=0.918, R^2=92.22%
Actual N:C Ratio
Predicted N:C Ratio
-0.5
0.0
0.5
1.0

**Table S1.** Mixed Effect Model Statistics for Bulk Metagenomic GC Content. A mixed effects model was used to estimate the difference in total average GC content of metagenomic reads between size fractions. The correlation length scale of the exponentially distributed random effect due to sample depth is reported along with estimates of the fixed effects for size fraction and sequencing depth on GC content.

| | **Average GC%** | | |
|---|---|---|---|
| *Predictors* | *Estimates* | *CI* | *p* |
| Intercept | 56.90 | 51.75 – 62.04 | **<0.001** |
| Free-Living Size Fraction | -5.32 | -6.51 – -4.13 | **<0.001** |
| Log10 Sequencing Depth | -1.56 | -2.29 – -0.83 | **<0.001** |
| **Random Effects** | | | |
| $\sigma^2$ | 3.50 | | |
| $\tau_{00\ depth}$ | 1.28 | | |
| ICC | 0.27 | | |
| $N_{depth}$ | 22 | | |
| Observations | 58 | | |
| Marginal $R^2$ / Conditional $R^2$ | 0.619 / 0.721 | | |

**Table S2.** Mixed Effect Model Statistics for Testing Significant Effect of Particle Fraction on Stoichiogenomic Parameters. Models were constructed for each stoichiometric parameter for each Domain separately. Estimates and standard errors for the difference in means between particle and free-living fractions are presented with p-value, as well as total model AIC and log-likelihood for the mixed effect model with a random depth effect. The likelihood ratio compares a model with the random depth effect to a model without a depth effect, and a p-value for the corresponding log-ratio test. For models with a significant depth effect (likelihood ratio test $p<0.05$), the Spearman rank correlation of the random depth effect with depth is presented with a p-value. A positive value means in general the random effect increases with increasing depth, a negative value means the effect decreases with increasing depth.

Mixed Effect Model Summaries

Test for Significant Effect of Size-Fraction on Stoichiogenomic Parameters

| Domain | Parameter | Effect | Covariate | Estimate | Standard Error | Degrees of Freedom | Test Statistic | p |
|---|---|---|---|---|---|---|---|---|
| Bacterial Models | | | | | | | | |
| Bacteria | Gene GC | Fixed | Intercept | 55.300000 | 1.850000 | 34 | 29.800 | 5.96e-26 |
| Bacteria | Gene GC | Fixed | Planktonic Fraction | -3.820000 | 0.481000 | 34 | -7.950 | 2.95e-09 |
| Bacteria | Gene GC | Fixed | Sequencing Depth (log10 reads) | -1.330000 | 0.248000 | 34 | -5.370 | 5.68e-06 |
| Bacteria | Gene GC | Random | Depth Random Effect | 1.810000 | NA | NA | NA | NA |
| Bacteria | Gene GC | Random | Intercept Random Effect | 1.910000 | NA | NA | NA | NA |
| Bacteria | Gene N:C Ratio | Fixed | Intercept | 0.123000 | 0.001640 | 34 | 75.000 | 2.32e-39 |
| Bacteria | Gene N:C Ratio | Fixed | Planktonic Fraction | -0.005160 | 0.000407 | 34 | -12.700 | 1.92e-14 |
| Bacteria | Gene N:C Ratio | Fixed | Sequencing Depth (log10 reads) | -0.000910 | 0.000225 | 34 | -4.050 | 2.81e-04 |
| Bacteria | Gene N:C Ratio | Random | Depth Random Effect | 0.001430 | NA | NA | NA | NA |
| Bacteria | Gene N:C Ratio | Random | Intercept Random Effect | 0.001380 | NA | NA | NA | NA |
| Bacteria | Gene #N per Side Chain | Fixed | Intercept | 0.360000 | 0.004920 | 34 | 73.200 | 5.35e-39 |
| Bacteria | Gene #N per Side Chain | Fixed | Planktonic Fraction | -0.013400 | 0.001020 | 34 | -13.200 | 6.62e-15 |
| Bacteria | Gene #N per Side Chain | Fixed | Sequencing Depth (log10 reads) | -0.001550 | 0.000687 | 34 | -2.250 | 3.08e-02 |
| Bacteria | Gene #N per Side Chain | Random | Depth Random Effect | 0.003820 | NA | NA | NA | NA |
| Bacteria | Gene #N per Side Chain | Random | Intercept Random Effect | 0.003130 | NA | NA | NA | NA |
| Archaeal Models | | | | | | | | |
| Archaea | Gene GC | Fixed | Intercept | 53.500000 | 2.760000 | 34 | 19.400 | 5.61e-20 |
| Archaea | Gene GC | Fixed | Planktonic Fraction | -4.580000 | 0.676000 | 34 | -6.770 | 8.86e-08 |
| Archaea | Gene GC | Fixed | Sequencing Depth (log10 reads) | -1.170000 | 0.397000 | 34 | -2.940 | 5.91e-03 |
| Archaea | Gene GC | Random | Depth Random Effect | 0.197000 | NA | NA | NA | NA |
| Archaea | Gene GC | Random | Intercept Random Effect | 2.270000 | NA | NA | NA | NA |
| Archaea | Gene N:C Ratio | Fixed | Intercept | 0.124000 | 0.002800 | 34 | 44.400 | 1.10e-31 |
| Archaea | Gene N:C Ratio | Fixed | Planktonic Fraction | -0.005390 | 0.000601 | 34 | -8.960 | 1.78e-10 |
| Archaea | Gene N:C Ratio | Fixed | Sequencing Depth (log10 reads) | -0.000646 | 0.000387 | 34 | -1.670 | 1.04e-01 |
| Archaea | Gene N:C Ratio | Random | Depth Random Effect | 0.002470 | NA | NA | NA | NA |
| Archaea | Gene N:C Ratio | Random | Intercept Random Effect | 0.001850 | NA | NA | NA | NA |
| Archaea | Gene #N per Side Chain | Fixed | Intercept | 0.360000 | 0.006530 | 34 | 55.100 | 7.90e-35 |
| Archaea | Gene #N per Side Chain | Fixed | Planktonic Fraction | -0.012800 | 0.001520 | 34 | -8.400 | 8.25e-10 |
| Archaea | Gene #N per Side Chain | Fixed | Sequencing Depth (log10 reads) | -0.001070 | 0.000921 | 34 | -1.170 | 2.52e-01 |
| Archaea | Gene #N per Side Chain | Random | Depth Random Effect | 0.003350 | NA | NA | NA | NA |
| Archaea | Gene #N per Side Chain | Random | Intercept Random Effect | 0.004860 | NA | NA | NA | NA |
| Viral Models | | | | | | | | |
| Virus | Gene GC | Fixed | Intercept | 39.700000 | 1.020000 | 34 | 38.900 | 9.14e-30 |
| Virus | Gene GC | Fixed | Planktonic Fraction | 0.384000 | 0.241000 | 34 | 1.590 | 1.21e-01 |
| Virus | Gene GC | Fixed | Sequencing Depth (log10 reads) | -0.465000 | 0.149000 | 34 | -3.120 | 3.67e-03 |
| Virus | Gene GC | Random | Depth Random Effect | 0.000040 | NA | NA | NA | NA |
| Virus | Gene GC | Random | Intercept Random Effect | 0.788000 | NA | NA | NA | NA |
| Virus | Gene N:C Ratio | Fixed | Intercept | 0.115000 | 0.001260 | 34 | 91.500 | 2.84e-42 |
| Virus | Gene N:C Ratio | Fixed | Planktonic Fraction | 0.000587 | 0.000319 | 34 | 1.840 | 7.43e-02 |
| Virus | Gene N:C Ratio | Fixed | Sequencing Depth (log10 reads) | -0.000152 | 0.000172 | 34 | -0.882 | 3.84e-01 |
| Virus | Gene N:C Ratio | Random | Depth Random Effect | 0.001010 | NA | NA | NA | NA |
| Virus | Gene N:C Ratio | Random | Intercept Random Effect | 0.001140 | NA | NA | NA | NA |
| Virus | Gene #N per Side Chain | Fixed | Intercept | 0.351000 | 0.004170 | 34 | 84.000 | 5.04e-41 |
| Virus | Gene #N per Side Chain | Fixed | Planktonic Fraction | 0.001160 | 0.001020 | 34 | 1.140 | 2.64e-01 |
| Virus | Gene #N per Side Chain | Fixed | Sequencing Depth (log10 reads) | 0.000504 | 0.000570 | 34 | 0.885 | 3.83e-01 |
| Virus | Gene #N per Side Chain | Random | Depth Random Effect | 0.003830 | NA | NA | NA | NA |
| Virus | Gene #N per Side Chain | Random | Intercept Random Effect | 0.003380 | NA | NA | NA | NA |

**Table S3.** Model Parameters for Multiresponse LASSO Regression Models. Table contains information for the parameter value in estimating protein N:C ratio for bacterial *rpoZ*, archaeal *ftsZ*, and viral Gp23, as well as the N-ARSC value of the amino acid.

| Amino Acid/Domain | Amino Acid | N:C Effect | N-ARSC of Amino Acid |
|---|---|---|---|
| Virus | | | |
| Arg___Virus | R | 1.394517e+00 | 3 |
| Gln___Virus | Q | 3.515107e-01 | 1 |
| His___Virus | H | 2.494210e-01 | 2 |
| Lys___Virus | K | 1.415429e-01 | 1 |
| Asn___Virus | N | 6.214760e-02 | 1 |
| Met___Virus | M | 5.508454e-02 | 0 |
| Trp___Virus | W | 3.890521e-03 | 1 |
| Asp___Virus | D | 2.825687e-03 | 0 |
| Ile___Virus | I | -9.346718e-03 | 0 |
| Leu___Virus | L | -3.779047e-02 | 0 |
| Ser___Virus | S | -4.949600e-02 | 0 |
| Gly___Virus | G | -1.548674e-01 | 0 |
| Glu___Virus | E | -2.932842e-01 | 0 |
| Val___Virus | V | -3.031260e-01 | 0 |
| Tyr___Virus | Y | -3.077628e-01 | 0 |
| Phe___Virus | F | -3.217233e-01 | 0 |
| Thr___Virus | T | -3.622502e-01 | 0 |
| Archaea | | | |
| Arg___Archaea | R | 1.264160e+00 | 3 |
| Asn___Archaea | N | 2.603076e-01 | 1 |
| His___Archaea | H | 2.405922e-01 | 2 |
| Lys___Archaea | K | 1.999981e-01 | 1 |
| Gln___Archaea | Q | 3.150754e-02 | 1 |
| Trp___Archaea | W | 2.150924e-02 | 1 |
| Gly___Archaea | G | -1.297567e-03 | 0 |
| Pro___Archaea | P | -1.179893e-02 | 0 |
| Thr___Archaea | T | -1.701073e-02 | 0 |
| Ser___Archaea | S | -4.764219e-02 | 0 |
| Ala___Archaea | A | -5.690299e-02 | 0 |
| Ile___Archaea | I | -6.632067e-02 | 0 |
| Val___Archaea | V | -6.692845e-02 | 0 |
| Asp___Archaea | D | -7.017555e-02 | 0 |
| Cys___Archaea | C | -7.470197e-02 | 0 |
| Phe___Archaea | F | -9.490948e-02 | 0 |
| Leu___Archaea | L | -9.609897e-02 | 0 |
| Met___Archaea | M | -1.184067e-01 | 0 |
| Glu___Archaea | E | -1.373760e-01 | 0 |
| Tyr___Archaea | Y | -2.318618e-01 | 0 |
| Bacteria | | | |
| Arg___Bacteria | R | 4.473871e-01 | 3 |
| Asn___Bacteria | N | 2.719465e-01 | 1 |

| Amino Acid/Domain | Amino Acid | N:C Effect | N-ARSC of Amino Acid |
|---|---|---|---|
| Lys___Bacteria | K | 1.583015e-01 | 1 |
| His___Bacteria | H | 5.907580e-02 | 2 |
| Gln___Bacteria | Q | 1.589507e-02 | 1 |
| Thr___Bacteria | T | -8.962773e-05 | 0 |
| Ala___Bacteria | A | -4.415793e-03 | 0 |
| Leu___Bacteria | L | -1.513929e-02 | 0 |
| Phe___Bacteria | F | -2.258591e-02 | 0 |
| Gly___Bacteria | G | -3.291845e-02 | 0 |
| Cys___Bacteria | C | -4.134283e-02 | 0 |
| Trp___Bacteria | W | -4.506462e-02 | 1 |
| Tyr___Bacteria | Y | -7.875854e-02 | 0 |
| Met___Bacteria | M | -9.827904e-02 | 0 |
| Pro___Bacteria | P | -1.917771e-01 | 0 |
| Asp___Bacteria | D | -2.948196e-01 | 0 |
| Glu___Bacteria | E | -5.088262e-01 | 0 |

**Table S4.** Model Statistics for Core Gene Mixed Effect Models. Mixed effect model statistics for model setups matching those presented in supplemental table 2, but for bacterial *rpoZ* archaeal *ftsZ* and viral Gp23 individually are presented.

Mixed Effect Model Summaries

Test for Significant Effect of Size-Fraction on Marker Genes

| Domain | Parameter | Effect | Covariate | Estimate | Standard Error | Degrees of Freedom | Test Statistic | p |
|---|---|---|---|---|---|---|---|---|
| Bacterial Models | | | | | | | | |
| Bacteria | rpoZ N:C | Fixed | Intercept | 1.21e-01 | 0.014300 | 33 | 8.430 | 9.64e-10 |
| Bacteria | rpoZ N:C | Fixed | Planktonic Fraction | -1.11e-02 | 0.002840 | 33 | -3.910 | 4.36e-04 |
| Bacteria | rpoZ N:C | Fixed | Sequencing Depth (log10 reads) | 3.71e-04 | 0.002030 | 33 | 0.183 | 8.56e-01 |
| Bacteria | rpoZ N:C | Random | Depth Random Effect | 3.69e-03 | NA | NA | NA | NA |
| Bacteria | rpoZ N:C | Random | Intercept Random Effect | 8.68e-03 | NA | NA | NA | NA |
| Archaeal Models | | | | | | | | |
| Archaea | ftsZ N:C | Fixed | Intercept | 9.56e-02 | 0.006400 | 34 | 14.900 | 1.66e-16 |
| Archaea | ftsZ N:C | Fixed | Sequencing Depth (log10 reads) | 1.91e-03 | 0.000946 | 34 | 2.010 | 5.20e-02 |
| Archaea | ftsZ N:C | Fixed | Planktonic Fraction | -8.14e-04 | 0.001420 | 34 | -0.572 | 5.71e-01 |
| Archaea | ftsZ N:C | Random | Depth Random Effect | 4.17e-07 | NA | NA | NA | NA |
| Archaea | ftsZ N:C | Random | Intercept Random Effect | 4.49e-03 | NA | NA | NA | NA |
| Viral Models | | | | | | | | |
| Virus | Gp23 N:C | Fixed | Intercept | 1.24e-01 | 0.006630 | 33 | 18.700 | 3.66e-19 |
| Virus | Gp23 N:C | Fixed | Planktonic Fraction | -3.73e-03 | 0.001530 | 33 | -2.440 | 2.02e-02 |
| Virus | Gp23 N:C | Fixed | Sequencing Depth (log10 reads) | -7.59e-04 | 0.000913 | 33 | -0.831 | 4.12e-01 |
| Virus | Gp23 N:C | Random | Depth Random Effect | 8.46e-04 | NA | NA | NA | NA |
| Virus | Gp23 N:C | Random | Intercept Random Effect | 5.84e-03 | NA | NA | NA | NA |
